# Transcriptome Alterations in Myotonic Dystrophy Frontal Cortex

**DOI:** 10.1101/2020.09.09.284505

**Authors:** Brittney A. Otero, Kiril Poukalov, Ryan P. Hildebrandt, Charles A. Thornton, Kenji Jinnai, Harutoshi Fujimura, Takashi Kimura, Katharine A. Hagerman, Jacinda B. Sampson, John W. Day, Eric T. Wang

**Affiliations:** Dept. of Molecular Genetics & Microbiology, Center for NeuroGenetics, Genetics Institute, University of Florida; Department of Neurology, University of Rochester Medical Center; Department of Neurology, National Hospital Organization Hyogo-Chuo Hospital; Department of Neurology, National Hospital Organization Toneyama Hospital; Department of Neurology, Hyogo College of Medicine; Department of Neurology, Stanford University

## Abstract

Myotonic dystrophy (dystrophia myotonica, DM) is caused by expanded CTG/CCTG microsatellite repeats, leading to multi-systemic symptoms in skeletal muscle, heart, gastrointestinal, endocrine, and central nervous systems (CNS), among others. For some patients, CNS issues can be as debilitating or more so than muscle symptoms; they include hypersomnolence, executive dysfunction, white matter atrophy, and neurofibrillary tangles. Although transcriptomes from DM type 1 (DM1) skeletal muscle have provided useful insights into pathomechanisms and biomarkers, limited studies of transcriptomes have been performed in the CNS. To elucidate underlying causes of CNS dysfunction in patients, we have generated and analyzed RNA-seq transcriptomes from the frontal cortex of 21 DM1 patients, 4 DM type 2 (DM2) patients, and 8 unaffected controls. One hundred and thirty high confidence splicing changes were identified, most occurring exclusively in the CNS and not in skeletal muscle or heart. Mis-spliced exons were found in neurotransmitter receptors, ion channels, and synaptic scaffolds, and we identified an alternative exon in GRIP1 that modulates association with kinesins. Splicing changes exhibited a gradient of severity correlating with CTG repeat length, as measured by optical mapping of individual DNA molecules. All individuals studied, including those with modest splicing defects, showed extreme somatic mosaicism, with a subset of alleles having >1000 CTGs. Analyses of gene expression changes showed up-regulation of genes transcribed in microglia and endothelial cells, suggesting neuroinflammation, and downregulation of genes transcribed in neurons. Gene expression of RNAs encoding proteins detectable in cerebrospinal fluid were also found to correlate with mis-splicing, with implications for CNS biomarkers of disease severity. These findings provide a framework for future mechanistic and therapeutic studies of CNS issues in DM.

## Introduction

Myotonic dystrophy (DM) is a multi-systemic, progressive disease caused by expanded CTG/ CCTG repeats in the 3’ UTR of the dystrophia myotonica protein kinase gene (DMPK, DM Type 1, DM1)^1^ or the first intron of the cellular nucleic acid binding protein gene (CNBP, DM Type 2, DM2)^2^, respectively. Both DM1 and DM2 are highly variable in age of onset, clinical features, and disease severity^3, 4^. Although DM is well studied in the context of peripheral symptoms such as myotonia and muscle weakness, central nervous system (CNS) symptoms are also common in DM and can contribute significantly to neurological impairment^3, 4^. These symptoms include hypersomnia, executive functioning deficits, memory deficits, and emotional disturbances^5, 6, 7^. Imaging studies show white and gray matter abnormalities in multiple brain regions, and ventricle enlargement (reviewed in^8^). However, molecular mechanisms driving these neurobiological changes remain largely unknown.

While there is strong evidence to support a pathomechanism in which MBNL proteins are functionally depleted in both peripheral and CNS tissues in DM^9, 10, 11, 12, 13, 14^, limited analyses have been performed to comprehensively identify transcriptome changes in the CNS across a broad subset of patients, assess the extent to which MBNLs are depleted, and address whether other RNA binding proteins are perturbed. RNA-seq has not been applied to CNS tissues in DM2, and the extent to which RBFOX proteins may modulate CNS pathogenesis in DM2^15^ is also unexplored. Furthermore, as opposed to skeletal muscle in which dysfunction is predominantly driven by myonuclei, the extent to which different cell types are affected in the CNS of DM-affected individuals is unknown. CUG foci have been observed in neurons, glia, and oligodendrocytes^9^, but the contribution of each cell type to pathology has not been extensively explored. Although transcriptome dysregulation has been extensively studied in peripheral DM1 tissues^16, 17, 18, 19^, the repertoire of mRNAs expressed in the CNS is distinct, and therefore a different set of exons may be mis-regulated in this tissue. Finally, although there are clear examples of how specific splicing events in peripheral tissues cause particular DM symptoms (Clcn1 and myotonia^20^, Bin1 and muscle weakness^21^, Scn5a and cardiac arrhythmias^19^), no clear examples exist in the CNS. To lay the groundwork required to answer some of these questions, here we generate and analyze transcriptomes derived from a set of post-mortem frontal cortex (FC) samples (Brodmann Area 10) from DM1 patients, DM2 patients, and unaffected controls. We identify high confidence mis-spliced exons that show a gradient of changes across patients and study how a mis-splicing event in GRIP1 may alter its efficacy as a synaptic adaptor. We analyze DM transcriptomes together with additional single cell transcriptome datasets^22^ to assess potential changes in cell type composition and determine which cell types are potentially responsible for changes in gene expression and splicing patterns.

Somatic instability is well established as a major driver of age of onset in DM1 and other repeat expansion diseases^23, 24, 25^, but technical challenges have precluded facile, direct assessment of full length repeat-containing alleles in the DM1 CNS. Furthermore, the CTG repeat lengths required to sequester sufficient MBNL to elicit robust mis-splicing have not been assessed in this tissue. New technologies such as long read sequencing provide some advantages over Southern blotting and small pool PCR^26^, but often still require amplification or sub-cloning, which can introduce some biases. Here, we apply a new optical mapping approach to size expanded CTG repeats at single molecule resolution in DM1; this approach allows for unbiased, amplification-free measurement of repeat lengths from genomic DNA^27, 28^.

Finally, while studies of transcriptomes can identify molecular events driving disease features, they may also suggest potential markers of disease status. Here, we study changes in the frontal cortex, but similar molecular changes may occur in other brain regions. The aggregate of these pathological processes may be reflected in the composition of cerebrospinal fluid (CSF), facilitating the development of accessible biomarkers to more precisely characterize disease severity and measure changes following therapeutic intervention. Indeed, CSF biomarkers have been developed for other neurological diseases such as Huntington’s Disease, C9ALS/FTD, and Alzheimer’s Disease^29, 30, 31, 32^; in parallel, investigators have used proteomics to broadly characterize proteins present in CSF. In DM1, splicing biomarkers correlate with muscle strength^33^ and potentially reflect the concentration of free MBNL in skeletal muscle^17^; subsequent approaches have been developed to profile RNAs present in urine^34^, potentially providing a non-invasive route to measure disease status. By integrating our transcriptome analyses with existing knowledge of proteins present in CSF, we lay critical groundwork to properly inform and interpret future biomarker discovery efforts.

## Results

### Splicing dysregulation in the DM1 frontal cortex exhibits a gradient of severity

We performed RNA-seq using RNA from post-mortem frontal cortex (Brodmann Area 10) of 21 DM1, 4 DM2, and 8 unaffected age- and sex-matched individuals (Fig. 1A, see Methods). All libraries satisfied typical quality metrics in FASTQC^35^ and were sequenced to a depth of at least 88 million reads to provide sufficient coverage for analyses of gene expression and alternative splicing. Percent spliced in (ψ) values were estimated by MISO^36^, and using a threshold of at least 20% change in mean ψ (p < 0.01, rank-sum test), 130 exons were identified to be differentially included between DM1 and unaffected individuals (Fig. 1B, C, Table S2). These consisted of exons in genes encoding key synaptic scaffolding proteins, cytoskeletal organization components, ion channels and neurotransmitter receptors, including some previously identified (MBNL2 exon 5, KCNMA1 exon 34, MAPT exon 3, CSNK1D exon 9), as well as novel candidates (GRIP1 exon 21, GABRG2 exon 10, DLGAP1 exon 20, and PALM exon 8). Some of these exons showed much greater variability in ψ among DM1 patients relative to unaffected individuals. A similar observation in muscle likely reflects differences in disease severity across patients, in part determined by the extent of MBNL sequestration^17^. To assess whether a similar phenomenon might exist in the CNS, we correlated ψ between pairs of exons across all unaffected and DM1 individuals (Fig. 1D). ψ for MBNL2 exon 5 correlated strongly with ψ for GABRG2 exon 10 (Pearson R = −0.89), and for GRIP1 exon 21 versus DLGAP1 exon 20 (Pearson R = 0.72). The distribution of R values for all pairwise comparisons of all 130 mis-spliced exons was strongly shifted to the right, as compared to a null distribution computed following shuffling of sample labels (Fig. 1E). These observations suggest that, similar to peripheral tissues, mis-splicing in the CNS may exhibit a spectrum of disease severity driven by a common upstream factor.

**Figure 1.**
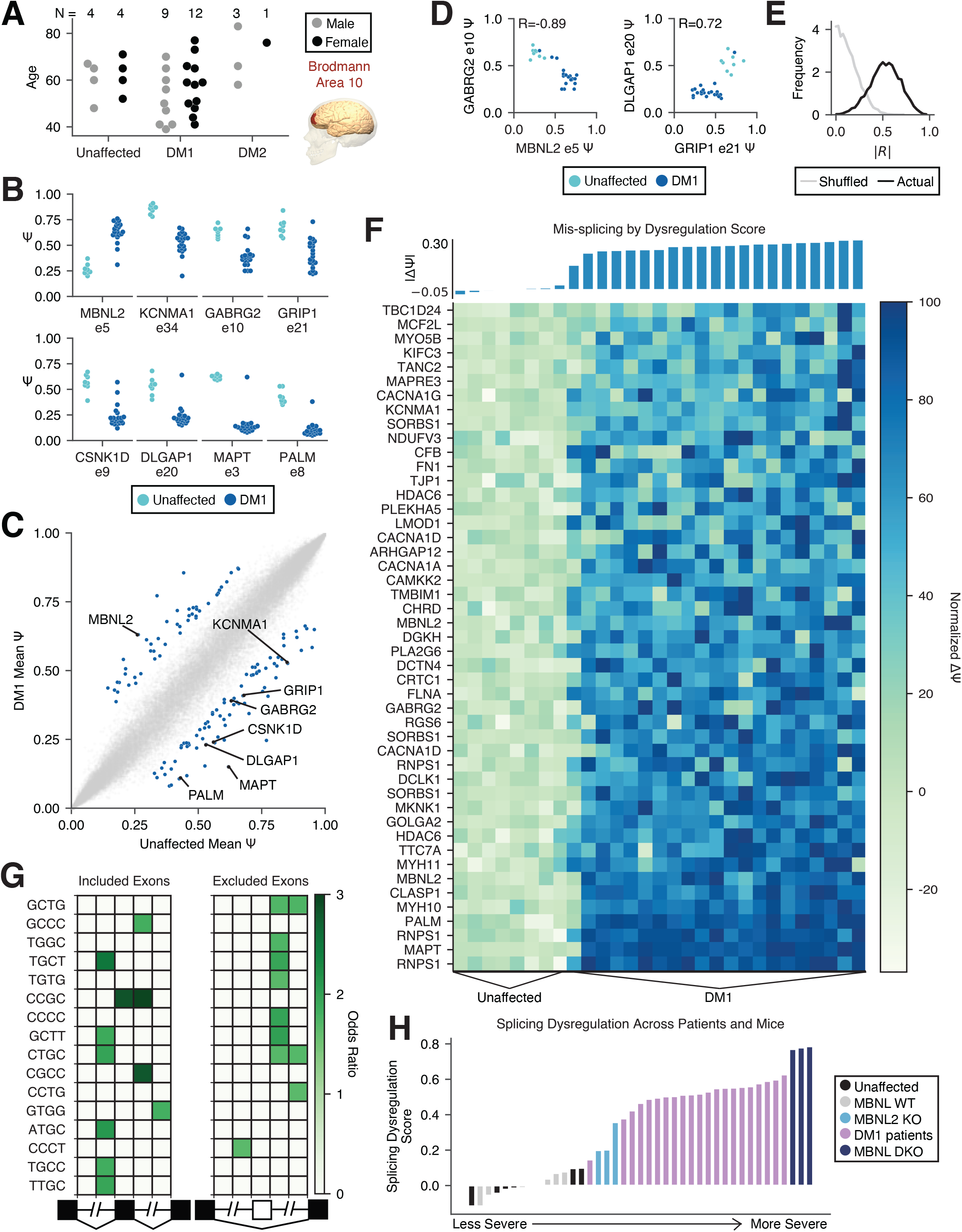
Mis-splicing changes across DM1 frontal cortex samples show a gradient of severity consistent with quantitative loss of MBNL. A) RNA-seq was performed on 21 DM1, 4 DM2 and 8 unaffected frontal cortex (Brodmann Area 10) samples across a range of ages. B) Percent spliced in (Psi, ψ) for specific exons in unaffected and DM1 frontal cortex samples. C) Scatter of mean ψ, unaffected versus DM1, for all exons measured; 130 mis-splicing events were detected as significantly regulated (|Δψ| > 0.2, p < 0.01 by rank-sum test), highlighted in blue. D) Scatter of ψ for MBNL2 exon 5 versus GABRG2 exon 10 and GRIP1 exon 21 versus DLGAP1 exon 20, across unaffected and DM1 individuals; Pearson correlation value is shown. E) Histogram of correlation values for all pairs of 130 significantly regulated exons (black). A similar histogram of correlation values for all pairs following shuffling of patient identities is also shown (gray, see Methods). F) Normalized ψ for 47 out of the 130 significantly regulated exons showing the least variation in ψ across unaffected frontal cortex (see Methods); individuals are sorted by “splicing dysregulation score” (see Methods), also shown above in bars. G) Heatmap showing enrichment of motifs around the 101 significantly regulated skipped exons relative to all other measured skipped exons. H) Total splicing dysregulation in all human and MBNL KO mouse samples considered, as computed using 74 orthologous exons significantly dysregulated (|Δψ| > 0.1, p < 0.05 by rank-sum test) in DM1 patients and MBNL DKO mice.

By visualizing a panel of 47 DM1-affected exons showing minimal variability among unaffected individuals, we observed that some events showed moderate dysregulation among some individuals while others showed much more widespread dysregulation, suggesting that some exons are more responsive to disease state than others (Fig. 1F). To more precisely quantitate overall mis-splicing, we computed a “Splicing Dysregulation Score” (Mean |Δψ|, see Methods), as performed similarly in peripheral tissues^17^ (Fig. 1F). This score did not correlate with age or sex (Fig. S1), and may be considered as a proxy for overall molecular perturbation, and it will remain to be seen in future studies whether this score correlates with disease symptoms.

### Mis-spliced exons in DM1 show MBNL motif signatures and suggest intermediate MBNL depletion

To identify *cis*-elements that may be associated with aberrant mis-splicing, we enumerated 4-mers in the regions flanking and within all mis-spliced skipped exons and compared to a set of CNS-expressed control skipped exons (see Methods). We observed a strong signature for MBNL motif (YGCY) enrichment, including TGCT and TGCC^37, 38^ (Fig. 1G). The pattern of enrichment also reflected expected functional binding patterns of MBNL, i.e. aberrantly included exons showed enriched binding sites within the upstream intron, and aberrantly excluded exons showed enriched binding sites within the downstream intron. These observations suggest that, similar to the periphery, MBNL sequestration is a major driver of mis-splicing in the DM1 CNS.

Several mouse models have been developed to study loss of MBNL function, including mice globally lacking MBNL2^11^ (MBNL2 KOs) and mice lacking MBNL1 constitutively and MBNL2 in neurons^12, 13^ (MBNL DKOs). To estimate the extent to which MBNL proteins are functionally depleted by expanded CUG repeats in these post-mortem samples, we calculated splicing dysregulation scores for WT mice, MBNL KO mice, and our post-mortem samples using a set of orthologous exons significantly mis-spliced in MBNL1/2 double KO mice and patients (see Methods). All patients except for one mild case showed splicing dysregulation between that observed in MBNL2 KO and MBNL1/2 double KO (Fig. 1H). In frontal cortex, at the RNA level, MBNL2 is ~2-fold more highly expressed than MBNL1, and if one were to assume that protein translation were similar for each MBNL, most individuals studied in this cohort show at least 66% depletion of MBNL.

### All assessed patients show CNS alleles with > 1000 CTGs and repeat lengths correlate with splicing dysregulation

To precisely measure repeat lengths at single molecule resolution, we used an optical mapping technique in which specific restriction sites are fluorescently labeled across the genome and imaged within DNA nanochannels^28^. We used specific labels in the vicinity of the DMPK locus to estimate CTG repeat lengths, where 1 micron spans approximately 2 kilobases (Fig. 2A, see Methods). We applied this approach to one unaffected control and 7 DM1 frontal cortex samples. While CTG repeat lengths in the unaffected control were estimated to be ~23 +/- 142 (std. deviation), a typical DM1 patient showed estimated lengths reaching ~5000. The distribution of lengths in the DM1 patients shifted rightwards such that the 50th, 75th, and 90th percentiles of repeat lengths were 84, 2306, 3910, respectively (Fig. 2B). Every single DM1 sample analyzed showed repeat lengths greater than 1000 CTG, and 86% of them showed alleles >4400 CTG. The 50th, 75th, and 90th percentiles of repeat length were correlated to the previously computed splicing dysregulation score (Fig. 2C); the strongest correlation was observed with the 90th percentile of repeat lengths, or the minimum length of the longest ~20% of all expansion-carrying alleles.

**Figure 2.**
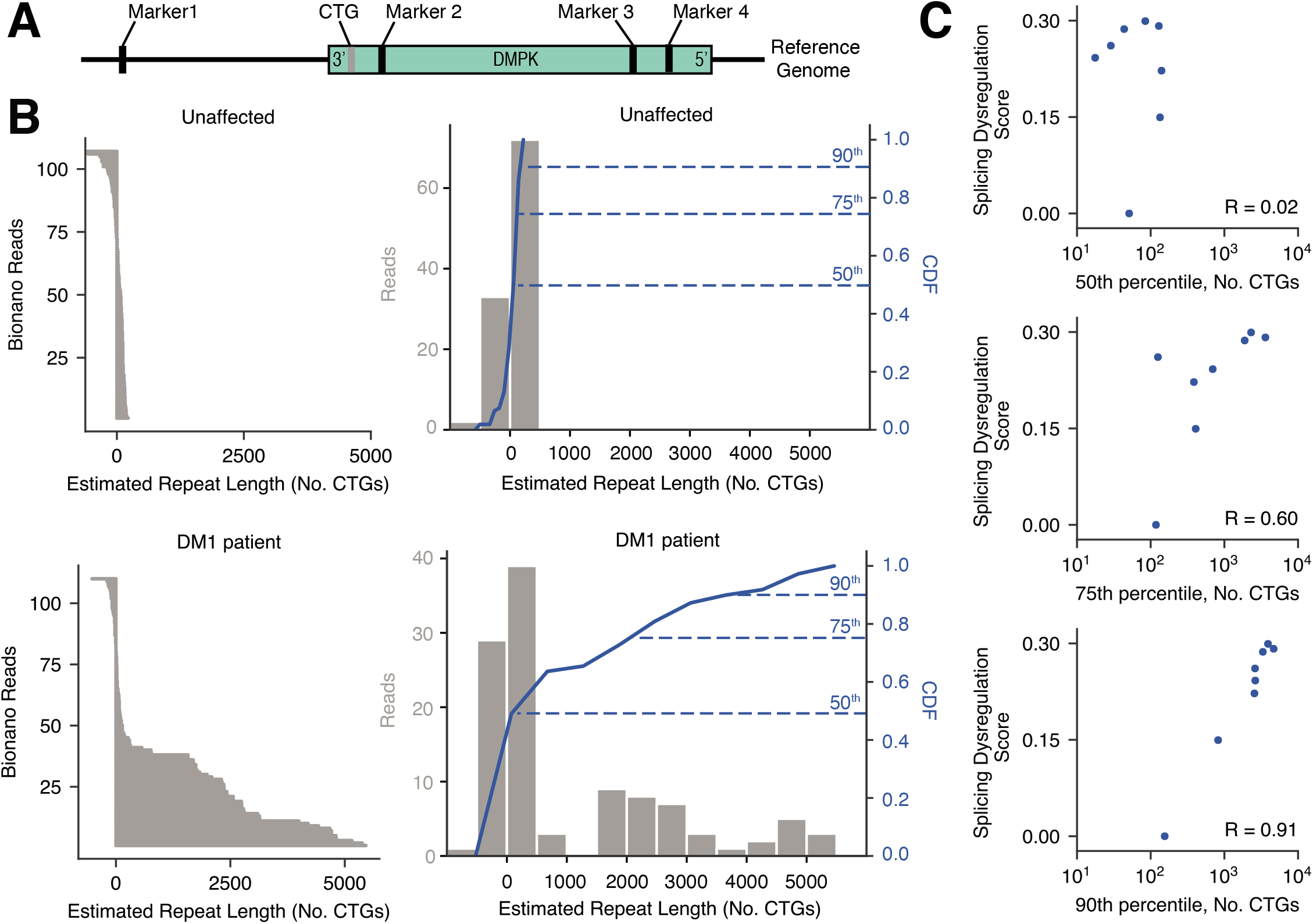
DM1 frontal cortex samples show large expansions, and the proportion of long expansions correlates with overall splicing dysregulation. A) DNA fragments from unaffected and DM1 frontal cortex samples were labeled by restriction nickases and subjected to optical mapping of the DMPK locus. B) Read density plots show distribution of estimated CTG repeat tract length and histograms of estimated CTG repeat tract length in each individual (gray bars), along with cumulative distribution function (CDF, blue line). The 50^th^, 75^th^ and 90^th^ percentiles of repeat lengths are indicated. C) Scatter of the 50^th^, 75^th^ and 90^th^ percentile of repeat lengths versus total splicing dysregulation across all unaffected and DM1 individuals for which repeat lengths were measured. Pearson correlation values are shown.

### Alternative splicing of the kinesin binding domain in GRIP1 is perturbed in DM1 and modulates association with KIF5A

As shown in Figure 1, multiple synaptic scaffolding proteins contain exons dysregulated in DM1. We focused on exon 21 of GRIP1 (Fig. 3A), a gene that plays important roles in localizing AMPA receptors to the synapse and regulating synaptic scaffolding through its 7 PDZ domains. Exon 21 lies within the kinesin binding domain (KBD), which was previously shown to mediate interactions with KIF5A, B, and C^39^; we therefore hypothesized that this exon might modulate the behavior of the KBD. To measure association of GRIP1 isoforms with KIF5A in cell culture, we generated fluorescent protein fusion constructs to study the behavior of the GRIP1 KBD with and without exon 21. We modified a previously developed centrosome recruitment assay^40^ and co-transfected these constructs with a BicD2-KIF5A tail flag-tagged construct into Neuro2A cells; BicD2 is a dynein adapter which directs the KIF5A C-terminal tail to the centrosome, along with its putative cargoes (Fig. 3B, see Methods). Association of FP-GRIP1 KBD with the KIF5A tail was quantified by measuring mean intensity at the centrosome relative to mean cytoplasmic intensity. Exon 21-containing GRIP1 KBD exhibited stronger centrosome recruitment as compared to exon 21-lacking GRIP1 KBD (Fig. 3C, see Methods). This increased association with KIF5A was quantitated in experiments in which GFP constructs with and without exon 21 were tested separately (Fig. 3D) or competitively, in which mCherry and GFP fusions for each isoform were co-transfected (Fig. 3E). In both cases, the GRIP1 KBD with exon 21 was more strongly recruited to the centrosome (p < 1e-25 for single transfections, p < 1e-26 for co-transfections), suggesting an important role for this exon to regulate KIF associations and synaptic recruitment of cargoes.

**Figure 3.**
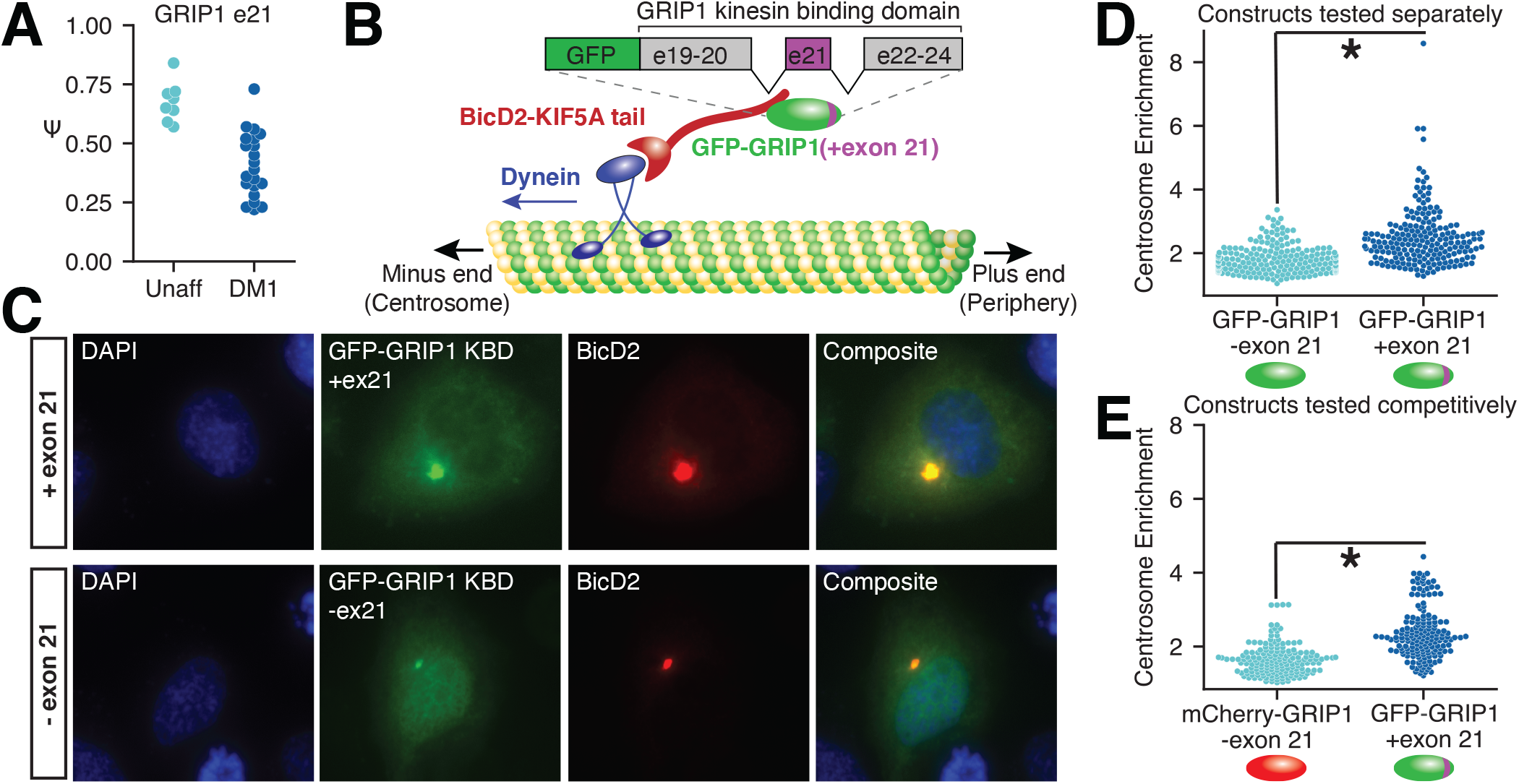
Mis-splicing of GRIP1 in DM1 may lead to changes in kinesin association. A) GRIP1 exon 21 is significantly mis-spliced in DM1 frontal cortex. B) Schematic of GRIP1 kinesin binding domain (KBD) and the centrosome recruitment assay. The C-terminal tail of KIF5A is used as a bait to recruit GFP-GRIP1 fusions to the centrosome. C) Representative images of the centrosome recruitment assay using GRIP1 KBD fluorescent fusion proteins, with and without exon 21, taken at 40x magnification. D) Quantitation of recruitment efficiency (mean signal at centrosome divided by mean cytoplasmic signal outside the centrosome) for each construct either transfected independently or E) in a competitive manner in which the construct with exon 21 was fused to GFP and the construct without exon 21 was fused to mCherry. Significance is shown by asterisk, p < 0.01 by rank-sum test.

### Splicing dysregulation in DM1 is largely tissue-specific

To explore potential overlaps in DM1 splicing dysregulation between frontal cortex and peripheral tissues, we analyzed previously published RNA-sequencing data from tibialis anterior (TA) and heart^16^. We identified exons mis-spliced in each tissue (|Δψ| ≥ 0.1, rank-sum p-value ≤ .01) and assessed 2- and 3-way overlaps. Interestingly, only ~10% of the exons dysregulated in the frontal cortex were also dysregulated in peripheral tissues; ~20% are simply not expressed in peripheral tissues, whereas ~70% are expressed in peripheral tissues but do not show the same dysregulation (Fig. 4A). Overlaps between any given pair of tissues revealed many dozens of shared dysregulated exons; some examples include NDUFV3 exon 3 (significantly regulated only in TA and FC) and DGKH exon 29 (significantly regulated only in heart and FC). The overlap in shared dysregulated exons among all three tissues was 25 shared exons, including MBNL1 exon 5, MBNL2 exon 5, and MAPT exon 3. While the number of total patients considered within each tissue can influence the total number of exons identified to be significantly dysregulated in any given tissue, we performed sub-sampling analyses to match patient group sizes, and found similar proportions of overlap (Fig. S2).

**Figure 4.**
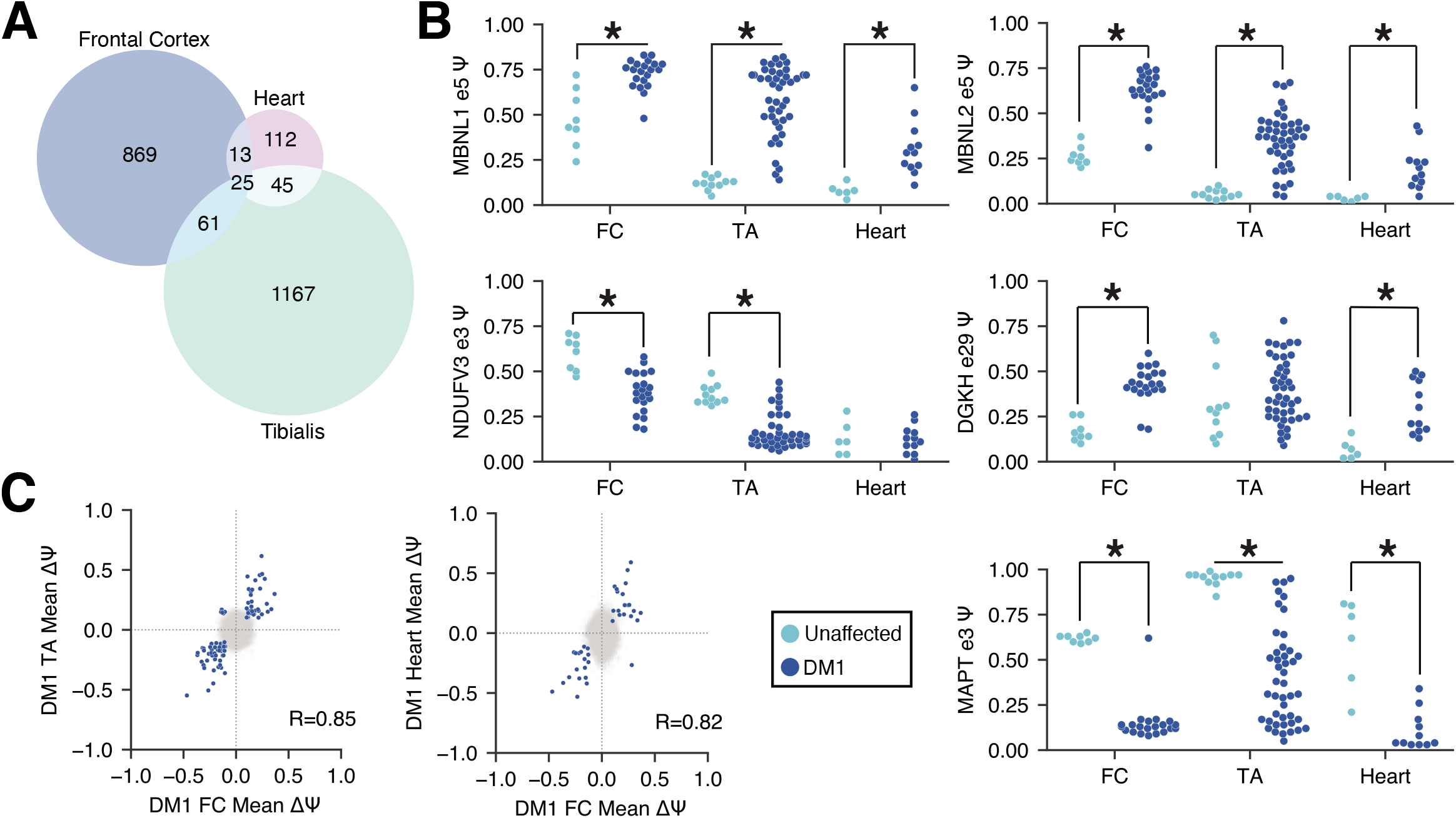
Across DM1 frontal cortex, skeletal muscle, and heart, most splicing events are tissue-specific. A) Venn diagram showing exons mis-spliced in 1, 2, or all 3 DM1 tissues. B) ψ for specific exons in the frontal cortex, skeletal muscle, and heart. Exons significantly misregulated in any given tissue are indicated with an asterisk. C) Scatter of ψ for each pair of tissues analyzed. Exons that are significantly regulated (|Δψ| > 0.1, p < 0.01 by rank-sum test) in both tissues are highlighted in blue; 86 are shared between the frontal cortex and skeletal muscle, and 38 between the frontal cortex and heart. Pearson correlations for blue points are shown.

Among the events that are shared between any pair of tissues, the direction of dysregulation was found to be conserved (Fig. 4C), as reflected by the strong correlations in Δψ between TA versus FC (Pearson R = 0.85), and heart versus FC (Pearson R = 0.82). These observations suggest similar mechanisms for splicing dysregulation across all tissues.

### Splicing alterations in DM2 are largely distinct but implicate secondary binding motifs for MBNL and RBFOX

DM1 and DM2 are caused by related repeat expansions (CTG and CCTG, respectively); they exhibit elements of shared CNS pathology, including problems with executive function and hypersomnolence, and potential elements of shared molecular pathology. Both diseases are thought to be driven by MBNL sequestration, but RBFOX may also play a role in DM2^15^. We profiled four DM2 frontal cortex samples, as there was limited availability of post-mortem tissue. Consistent with previous observations in muscle and blood cells, intron 1 of CNBP was retained in all DM2 samples, but was efficiently spliced in all DM1 and unaffected samples^41, 16^ (Fig. S3). We compared mis-splicing in DM2 to that observed in DM1 and identified exons exclusively regulated in one disease or shared by both (|Δψ| ≥ 0.1, rank-sum p-value ≤ 0.01, Table S4). Approximately ~28% of exons dysregulated in DM2 were also dysregulated in DM1 (Fig. 5A), and generally changed in the same direction (Fig. 5B, shown in teal). Many exons unique to DM2 (Fig. 5B, shown in purple) trended towards regulation in DM1 but did not reach significance. To estimate the extent to which our sample size for DM2 might limit the power to detect dysregulated exons in DM2, we repeatedly sub-sampled 4 out of 21 DM1 samples and re-estimated the overlap between DM2- and DM1-regulated events. We estimate that with the 4 DM2 samples analyzed, we capture 25-50% of the total repertoire of altered splicing that may exist in DM2 (Fig. S3, see Methods).

**Figure 5.**
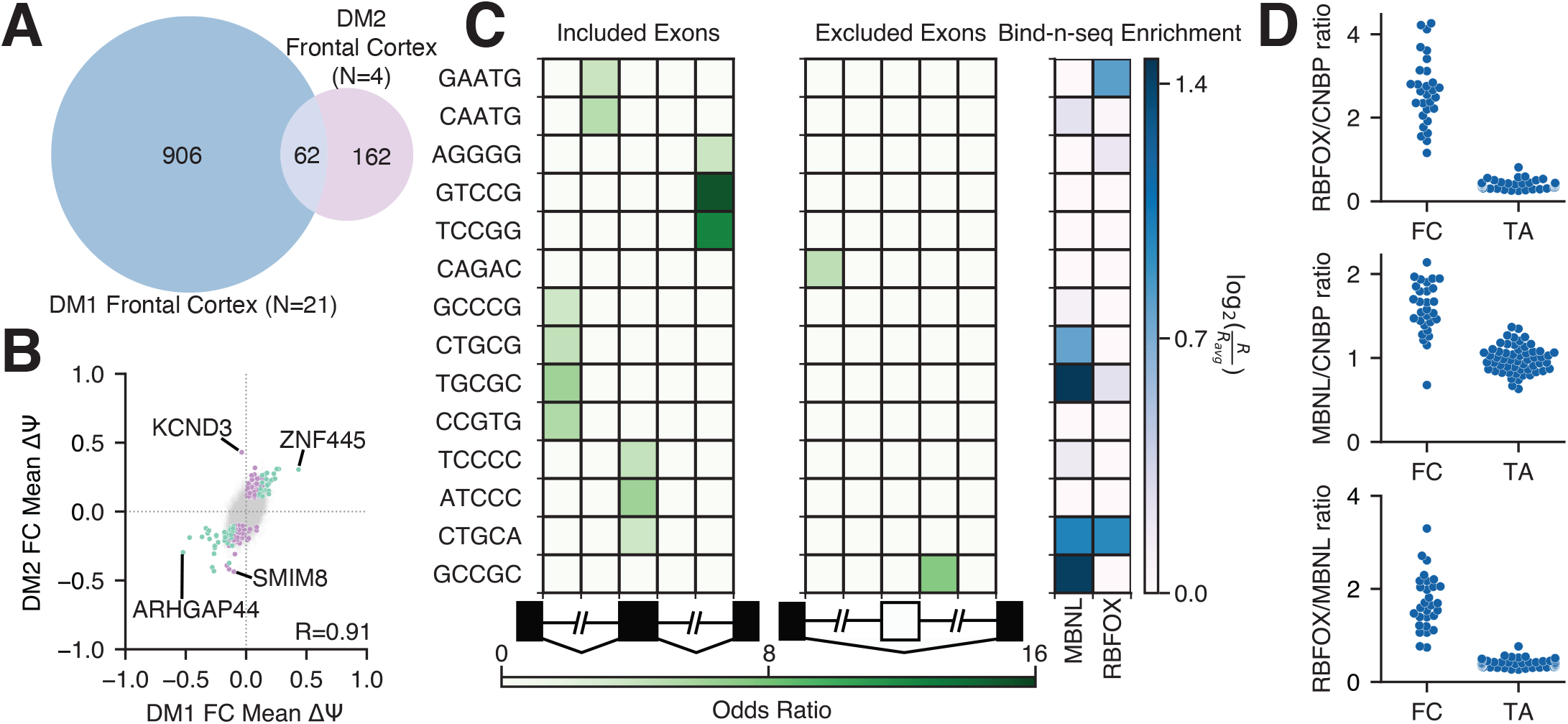
The extent of mis-splicing shared between DM1 and DM2 frontal cortex is limited. A) Venn diagram showing exons mis-spliced (|Δψ| > 0.1, p < 0.01 by rank-sum test) in DM1 and DM2 frontal cortex, or both. B) Scatter of Δψ, DM2 versus DM1. 62 exons were identified to be significantly regulated in both DM1 and DM2 (|Δψ| > 0.1, p < 0.01 by rank-sum test) are highlighted in teal, 162 exons were identified to be significantly regulated uniquely in DM2 (|Δψ| > 0.1, p < 0.01 by rank-sum test) are highlighted in purple. DM1-specific events have been omitted for clarity. Pearson correlation for shared (teal) events is shown. C) Heatmap showing enrichment of motifs around 35 DM2-regulated skipped exons relative to all other measured skipped exons. Bind-N-Seq enrichment values for MBNL and RBFOX are also shown; enrichments were derived from experiments using 1080 nM MBNL1 or 1100 nM RBFOX2. D) TPM ratios of total RBFOX (RBFOX1, 2, 3) versus CNBP, total MBNL (MBNL1, 2, 3) versus CNBP and total RBFOX versus total MBNL are shown across frontal cortex and skeletal muscle; note the high concentration of RBFOX in frontal cortex relative to skeletal muscle.

To identify *cis*-elements that may play a role in regulating DM2-specific mis-splicing events, we enumerated 5-mers in regions flanking and within all significantly regulated skipped exons and compared to a set of CNS-expressed control skipped exons (see Methods). In contrast to analyses of exons differentially regulated in DM1, enrichment of canonical MBNL and RBFOX motifs was not readily apparent (Fig. 5C, left panel). The absence of canonical YGCY and UGCAU/GCAUG motifs for MBNL and RBFOX, respectively, motivated us to examine the expression levels of MBNL, RBFOX, and CNBP genes in TA and frontal cortex, to assess whether the stoichiometry of each player might provide additional insights (Fig. 5D). In all comparisons, we found that the ratio of gene expression for MBNL (1, 2 and 3) and/or RBFOX (1, 2 and 3) versus CNBP was much higher in frontal cortex relative to TA, suggesting a greater buffering capacity (i.e. resistance to depletion) in frontal cortex. In contrast, the ratio of DMPK to MBNL is similar in frontal cortex relative to TA, consistent with similar signatures of MBNL motif enrichment around exons mis-spliced in DM1 frontal cortex and TA^16^.

Analyses of *in vitro* binding preferences for RNA binding proteins (RBPs) indicate that the concentration of the RBP can influence the extent to which optimal versus sub-optimal motifs are bound^37, 42^. That is, the “best” motifs are bound when RBP concentration is limiting, but “sub-optimal” motifs are also bound when RBP concentration is high. Indeed, upon analyzing Bind-N-Seq data for MBNL1 and RBFOX2 at elevated protein concentrations^37^ (1080 nM for MBNL1 and 1100 nM for RBFOX2), we observed positive binding enrichments for both MBNL and RBFOX among DM2-enriched motifs (Fig. 5C, right panel). We also assessed whether exons exclusively regulated in DM2 and not in DM1 showed enrichment for sub-optimal RBFOX binding sites, but could not detect any effects reaching statistical significance (data not shown).

### Gene expression changes suggest neuroinflammation, are cell type-specific, correlate with splicing dysregulation, and reveal potential biomarkers

Using Kallisto^43^ and Sleuth^44^, we estimated gene expression changes (Table S4) for all DM1 and unaffected frontal cortex samples. Two hundred thirty-five genes were significantly up-regulated (q < 0.01 and |fold-change| ≥ 1) in the DM1 frontal cortex, and 145 genes were significantly down-regulated. Gene Ontology analyses (Fig. 6A) showed that up-regulated genes were enriched in adaptive immune response, cell regulation of leukocyte proliferation, and inflammatory response. Down-regulated genes were enriched in actin cytoskeleton organization, regulation of cation transmembrane transport, and regulation of synaptic plasticity. Because the frontal cortex contains a variety of cell types, we investigated whether there might be changes in cell type composition between patients and unaffected controls. Using published single cell RNA-seq data^22^, we built a Bayesian inference approach to estimate the proportion of neurons, endothelial cells, astrocytes, oligodendrocytes, microglia, and oligodendrocyte precursor cells within each sample. This model was able to detect frank neuronal cell loss in the context of Alzheimer’s Disease^45^ (Fig. S4) but did not show neuronal cell loss in DM1 (Fig. 6B). However, we did observe a signature for increased microglial composition among patients, consistent with an inflammatory response (Fig. 6B) and gliosis that has been previously observed^46^. We classified up- and down-regulated genes by their cell type of origin (genes expressed similarly across multiple cell types were omitted from this analysis; see Methods) and observed a strong signature for neuronal genes being down-regulated, as well as a strong signature for endothelial and microglial genes being up-regulated (Fig 6C, left panel). In contrast, splicing changes were inferred to be present in all cell types studied (Fig. 6C, right panel). The ratio of DMPK to MBNL (1, 2 and 3) was highest in endothelial cells, astrocytes, and neurons, perhaps providing conditions most conducive to strong MBNL sequestration in those cell types (Fig. 6D).

**Figure 6.**
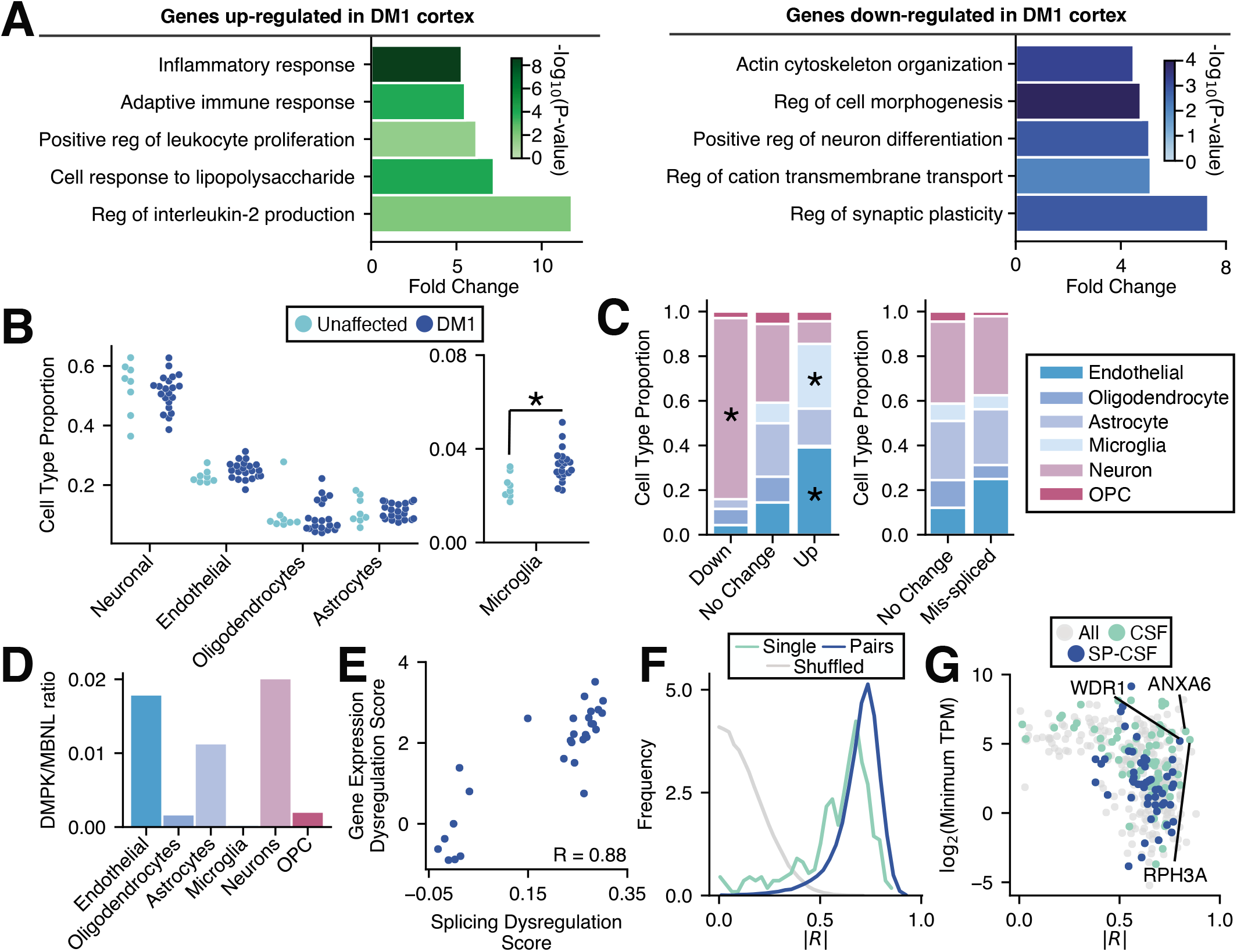
Analysis of gene expression changes reveals neuroinflammation and potential biomarkers. A) Gene Ontology analysis of genes up- (green) and down-regulated (blue) in DM1 versus unaffected frontal cortex (top five categories shown for each, selected by fold change). B) The proportion of neurons, microglia, endothelial cells, oligodendrocytes, and astrocytes in each frontal cortex sample was estimated by Bayesian inference using published transcriptome profiles (see Methods) and plotted. Significance is shown by asterisk. C) The proportion of up-, non-, and down-regulated genes derived from specific cell types is shown (left panel). The proportion of mis-spliced and unaffected genes derived from specific cell types is also shown (right panel). Cell type specificity for genes was determined using publicly available single cell sequencing data (see Methods). Significance is shown by asterisk. D) TPM ratios for DMPK versus total MBNL (MBNL1, 2, 3) across various CNS cell types. E) Scatter of splicing dysregulation score versus gene expression dysregulation score (see Methods). Pearson correlation is shown. F) Pearson correlations between splicing dysregulation score and log2(TPM) for each dysregulated gene were computed and plotted as a histogram (teal); samples were shuffled and correlations were recomputed and plotted (gray). A similar score was also computed between splicing dysregulation and all pairs of genes (blue, see Methods). Absolute values for all correlations were used for plotting. G) Scatter of the single gene correlations computed in (F) versus log2(TPM) for those genes. Genes encoding proteins detectable in CSF, and those additionally found to have signal sequences (SignalP, see Methods) are highlighted in teal or blue, respectively.

We next investigated whether gene expression changes correlate with splicing dysregulation across DM1 and unaffected controls. We developed a “Gene Expression Dysregulation Score” which summarized the total change in gene expression among up- and down-regulated genes (see Methods). These scores were then correlated with previously calculated splicing dysregulation scores, revealing a strong correlation (Pearson R = 0.88, Fig. 6E). To investigate whether a smaller subset of genes could be as informative as assessment of all dysregulated genes, we correlated individual and pairs of gene expression changes to splicing (Fig. 6F) and compared to correlations of shuffled data. Many gene expression changes showed correlations > 0.8 and may be strong candidates for markers of pathology. For purposes of clinical studies or trials, however, a marker would ideally be accessible via cerebrospinal fluid (CSF). We overlapped our frontal cortex gene expression changes with proteomics studies of CSF^47^ and identified genes that 1) encode proteins detectable in the CSF (with or without canonical signal sequences, determined by SignalP^48^), 2) show strong correlations with splicing dysregulation and 3) are expressed at reasonable levels in the CNS to maximize sensitivity of detection (Fig. 6G). Several candidates are highlighted; WDR1 is up-regulated in the DM1 frontal cortex, and ANXA6 and RPH3A are down-regulated. WDR1 is involved in disassembly of actin filaments and cell migration^49^, ANXA6 is involved in vesicle fusion^50^, and RPH3A plays essential roles in synaptic vesicle release^51^. These gene expression changes may also be reflected at the RNA or protein level in DM CSF, providing an accessible route by which to measure CNS disease status in future studies.

## Discussion

In this study, we have generated and characterized transcriptomes derived from DM1 and DM2 frontal cortex and associated controls. We identified a high confidence set of splicing events whose dysregulation correlates with CTG repeat length as measured by optical mapping, an amplification-free approach to sizing repeats at a single molecule level. While previous analyses of progenitor allele lengths have observed correlations to age of onset and disease severity^52^, a correlation between actual repeat lengths and extent of mis-splicing in the same tissue has never been measured. Although we did not have phenotypic information about each individual, these observations suggest that the extent of mis-splicing and repeat length may also underlie severity of disease symptoms. Alterations in the splicing patterns of many neurotransmitter receptors and synaptic scaffolding proteins were identified, revealing candidates that may play important roles in the neurobiology of DM1 symptoms. Notably, mis-splicing of some exons tended to enhance the activity of inhibitory machinery and dampen the excitatory machinery. For example, skipping of GABRG2 exon 9 is associated with increased sleep times in mice treated with benzodiazepines^53^, and skipping of exon 8 of PALM, a developmentally regulated exon, is associated with reduced filopodia formation, dendritic spine maturation and AMPA receptor recruitment^54^. Exon skipping within the kinesin binding domain of GRIP1^39^, an important synaptic adaptor for AMPA receptors, may decrease the efficiency by which AMPA receptors are recruited and stabilized at synapses.

Analyses of *cis*-elements flanking dysregulated exons revealed that MBNL-bound motifs were the predominant signature, suggesting that functional depletion of MBNL is a major driver of splicing changes in the DM1 frontal cortex. Consistent with these observations, orthologous alternative exons in MBNL2 KO and MBNL1/2 double KO mice showed changes in splicing similar to DM1, with the severity of mis-splicing in virtually all DM1 patients showing intermediate depletion, e.g. between single MBNL2 KO and double MBNL1/2 KO. When considered together with estimates of repeat lengths we obtained using optical mapping, these observations suggest that CTG tracts of over 1000 repeat units, and perhaps up to 4000, are required to achieve this level of sequestration. This could explain why some animal and cell-based DM1 models do not recapitulate the full extent of mis-splicing observed in human tissue. Notably, correlations between mis-splicing and repeat lengths were strongest when considering the longest 20% of disease alleles (the longest 10% of all alleles), suggesting that somatic instability plays a critical role in disease onset and severity and that potentially, a subset of cells are responsible for the most severe molecular signatures observed. The expansion process may take decades, consistent with some DM1 symptoms only manifesting in adult or late adult life. Given that full depletion of MBNL leads to nuclear export of expanded DMPK^55^, it is possible that cells with extreme repeat lengths contain CUG-derived repeat-associated non-AUG translation peptides, invoking additional pathological mechanisms. Future work should focus on which cell types harbor these extremely large alleles, and the extent to which those cells contribute to the bulk transcriptome signatures we observe here. While we have focused on the frontal cortex in this study, the clinical presentation of DM and imaging studies indicate abnormalities in multiple brain regions. Future studies should similarly examine transcriptome changes and repeat length distributions in those regions so that molecular features can be better linked to particular symptoms. Of particular interest may be the hippocampus and amygdala, because these regions show anatomical differences in DM1 and are known to play roles in memory processing and emotional experiences^56^.

Surprisingly few transcripts showed mis-splicing across DM1 TA, heart, and FC. Even when exons are expressed in all 3 tissues, many show mis-splicing in only 2 out of 3 tissues. Tissuespecific regulation of RNA binding proteins or *trans*-factor environments may be responsible for these differences. While a greater number of DM2 frontal cortex samples will be needed to fully characterize transcriptome changes in DM2, we found limited overlap in mis-splicing between DM1 and DM2. Although shared events were generally dysregulated in the same direction in both DM1 and DM2, there was also a substantial subset of events dysregulated in only DM1 or only DM2. These observations may differ from that in peripheral tissues, but gene expression levels of RBFOX family members relative to MBNL or CNBP are much greater in the CNS as compared to skeletal muscle. RBFOX proteins have been proposed to buffer MBNL sequestration^15^ and this phenomenon may play a more substantial role in the CNS, where RBFOX proteins may be in greater stoichiometric excess than in muscle. Finally, differences in cell type-specific expression of DMPK, CNBP, MBNL, and RBFOX may also play a role in producing distinct transcriptome signatures of each disease subtype.

By implementing a Bayesian inference approach to infer cell type composition using published single cell RNA-seq data, we found no significant neuronal cell loss in the frontal cortex samples we studied. However, there was a significant increase in microglia. Down-regulated genes tended to be those expressed preferentially in neurons, and up-regulated genes tended to be those expressed preferentially in endothelial and microglial cells. These observations suggest a neuroinflammatory response accompanied by neuronal dysfunction but not overt neurodegeneration, at least in this brain region. The complexity of cell type composition in the CNS concurrent with somatic instability prompts future studies to directly assess variability of transcriptome alterations at a single cell level, to separate those cells driving disease versus those responding to paracrine signals. Overall, gene expression changes correlate strongly with mis-splicing, providing an opportunity to develop gene expression or protein-based biomarkers of disease severity. Indeed, overlapping observed gene expression changes with published CSF proteomics datasets suggest strong candidates for further evaluation. Notably, RPH3A plays roles at both the pre- and post-synapse; it interacts with RAB3A^57^ and SNAP25^58^ to regulate synaptic vesicle trafficking at the pre-synapse, and interacts with PSD-95 to stabilize NMDA receptors at the post-synapse^59^. Related to this pathway, RAB3A protein has been observed to be elevated in a CNS mouse model of DM as well as in DM1 frontal cortex^60^. Finally, RPH3A has also been observed to be downregulated in HD patients^61^. Therefore, in addition to holding potential utility as biomarkers, these candidates may also inform us about pathways perturbed specifically in DM, or broadly across neurological diseases.

## Data Availability

All RNA-seq data is available in GEO (accession number GSE157428).

All supplemental tables can be found at: http://ericwanglab.com/otero_fc.php

## Acknowledgments

We thank Kendra McKee for assistance with RNA-seq library preparation and Hailey Olafson for assistance downloading and demultiplexing fastq files. We thank Lance Denes for assistance cloning the pcDNA-BicD2 kinesin plasmids. We thank Gary Bassell for bringing to our attention the kinesin centrosome recruitment assay developed by Gary Banker, and providing a BicD2 template plasmid.

## Materials and Methods

### Tissue and RNA collection, RNA-seq library construction

Autopsy samples were obtained from Stanford University, The Research Resource Network Japan, University of Rochester Medical Center and the NIH Biobank. Tissue samples were homogenized in TRIzol using the Omni bead ruptor followed by the Direct-zol RNA Miniprep kit, with DNAse I treatment. RNA quality and abundance were assessed using fragment analysis (Agilent/Advanced Analytical). Samples with an RQN >4 were further processed. RNA-seq libraries were constructed using the NEBNext Ultra II Directional RNA library prep kit for Illumina, using ribosomal RNA depletion followed by strand-specific RNA-seq preparation. Samples were amplified with PCR for 9-11 cycles and sequenced using the Illumina NextSeq 500 v2 with 75 nucleotide paired end reads. Samples were sequenced across multiple runs to account for possible batch effects, and reads were pooled so that all samples had at least 88 million reads.

### Read mapping, gene expression quantitation and isoform quantitation

Upon passing FASTQC metrics for quality control^35^, reads were mapped to the hg19 genome using STAR^62^. Gene expression was quantified using Kallisto^43^, with hg19 Refseq tables as a reference. Differentially regulated genes were identified using Sleuth^44^. Isoform percent spliced in (PSI, Ψ) values were quantified by MISO^36^ using hg19 v2.0 MISO annotations (http://genes.mit.edu/burgelab/miso/annotations/ver2/miso_annotations_hg19_v2.zip).

### Identification of dysregulated exons and calculation of splicing dysregulation score

Splicing events were assessed for significant dysregulation across individuals by rank-sum test. One hundred thirty DM1 events and 59 DM2 events were found to meet criteria of |Δψ| > 0.2, p < 0.01 (rank-sum test). To assess a false discovery rate, psi values were shuffled among individuals for each event, and the rank-sum test was performed again. After shuffling, 6 out of 130 events remained significant, so we estimate a false discovery rate of <5%. Of the 130 significantly regulated exons in the DM1 frontal cortex, those with a psi range <0.25 among unaffected samples were chosen and shown in the heatmap in Fig. 1F. Patients were ordered along the x-axis by their splicing dysregulation score. This score was derived from the average magnitude of delta psi across all 130 significantly regulated exons, compared to the mean psi of the unaffected samples. The scores themselves are visualized in the corresponding bar graph above the heatmap.

### Splicing event cross-correlations

Cross-correlations were computed for psi values across individuals, between every pair of differentially regulated exons. The histogram of Pearson correlation values is shown in Fig. 1E in black. To compute a null distribution, psi values were shuffled among individuals, and Pearson correlations were re-computed between all pairs. The histogram of Pearson correlations for shuffled values is shown in gray.

### Motif Analysis

Of the 130 significantly regulated splicing events in the DM1 frontal cortex, 101 skipped exon events (31 aberrantly included exons and 70 aberrant excluded exons) were used to analyze motif signatures. Sequences from 5 distinct regions were assessed for each event: 250 bases downstream of the upstream exon, 250 bases upstream of the skipped exon, the skipped exon itself, 250 bases downstream of the skipped exon, and 250 bases upstream of the downstream exon. Four-mers within these regions were enumerated and compared to the same regions around all other skipped exon events observed in our dataset. Significance for 4mer enrichment was determined using a binomial test with Bonferroni correction. A similar analysis was performed for the top 59 events in DM2 frontal cortex, 35 of which were skipped exon events (17 aberrantly included exons and 18 aberrantly excluded exons). To provide additional resolution with respect to motif sequences, which facilitated comparison to 5mer Bind-N-Seq data^37^, 5mers were enumerated along each sequence from these regions, rather than 4mers.

### Analysis of exons conserved between human and mouse

Previously published RNA-seq datasets of MBNL2 KO^11^ and MBNL1/2 Nestin-cre DKO^13^ mice were analyzed. MISO was used to calculate PSI values across all captured events and significance was calculated by rank-sum test. Events were filtered by |Δψ| > 0.1, p < 0.05 and intersected with significantly mis-spliced events in the DM1 patients using human-mouse orthologous exons as defined by Ensembl. Seventy-nine exons were found to be dysregulated in both human DM1 patients and MBNL DKO mice, and 39 exons were found to be dysregulated in both human DM1 patients and MBNL2 KO mice. Of the 79 exons in both human DM1 and MBNL DKO, 77 were observed in all 3 datasets and used to calculate a splicing dysregulation score as defined above. Samples were ordered along the x-axis according to this score in Fig. 1H.

### Assessment of cell type composition by Bayesian Inference

Single-cell RNA-sequencing of human adult temporal lobe^22^ was analyzed to obtain cell typespecific markers in neurons, endothelial cells, microglia, astrocytes, oligodendrocytes, and oligodendrocyte precursor cells (OPCs). The top 50 enriched genes in each cell type were compiled and of these 300, 220 were expressed in our dataset. Using these cell type markers, a Bayesian inference model was built using Pymc3^63^ to estimate relative proportions of all cell types across all samples. We estimated the proportion of each cell type by implementing a Bayesian framework in which p(cell type | expression data) ∝ p(expression data | cell type) * p(cell type). The priors for each cell type were initialized as Dirichlets, and the sum of all cell type proportions was constrained to 100%. Reported cell type proportions are derived from the mean of the inferred posterior distributions. The estimated proportion of OPCs was <2.5% and was eliminated from the model; final cell type proportions are reported using a model that incorporates only neurons, endothelial cells, microglia, astrocytes, and oligodendrocytes and uses 184 cell type marker genes.

### Assessment of cell type-specific gene expression and splicing changes

Cell type specificity of genes was calculated by dividing the recorded expression of each gene in a given cell type by the expected expression of that gene if that gene were equally expressed across all cell types. We designated genes with scores >3 as specific. The number of up- and down-regulated genes (and genes that do not change) was enumerated. A Fisher exact test was used to determine whether particular cell types showed enrichment of differentially regulated genes versus genes not showing changes. A similar analysis was performed with differentially regulated exons versus all exons captured in the dataset not showing regulation.

### Gene ontology analysis

Gene Ontology analysis was performed on all significantly up- and down-regulated genes (235 and 145 genes, respectively), with all expressed genes in the dataset used as the background, using http://geneontology.org/. Categories were collapsed using Revigo^64^ and the top 5 categories (sorted by enrichment) are shown.

### Gene expression dysregulation score, correlations, and overlap with CSF proteomics and SignalP

The gene expression dysregulation score was calculated using the significantly up- and down-regulated genes; it was computed as the average magnitude of logged gene expression changes relative to the mean log2(TPM) of all unaffected samples. log2(TPM) for all up- and down-regulated genes were correlated to the splicing dysregulation score across all individuals (Fig. 6F). Pearson correlations were calculated and shown on the histogram in green. Additionally, scores were derived for all possible pairs of genes among significantly up- and down-regulated genes; the score for pairs of genes going in the same direction was defined as the sum of log(TPM) and the score for pairs of genes going in opposite directions was defined as the difference of log(TPM). Pearson correlations for these scores relative to splicing dysregulation score were calculated and shown on the histogram in blue. To obtain a null distribution for correlations, expression values for all up- and down-regulated genes were shuffled and compared again to splicing dysregulation; Pearson correlations were calculated and shown on the histogram in gray. Regulated genes were intersected with genes detected in CSF by proteomics^47^ to determine whether any genes would be strong biomarker candidates (Fig. 6G), shown in teal. This subset of genes was also scored by SignalP version 5^48^ to identify signal peptides, shown in blue.

### BioNano Saphyr sample preparation and mapping

For each sample, 10 mg brain tissue was homogenized, embedded in agarose plugs and incubated with lysis buffer and Proteinase K at 50°C overnight. RNase A was added and incubated for 1 hour at 37°C. Agarose was digested with agarase and the purified DNA was subjected to drop dialysis for 4 hours and quantified by the Qubit Broad Range DNA assay. Purified DNA was labeled using the Direct Label Enzyme (DLE) method (BioNano Genomics). A total of 750ng of DNA was labeled using the DLE-1 kit following the manufacturer’s instructions and then treated with proteinase K. The DNA was stained with YOYO-1 according to the DLE-1 kit instructions and homogenized by HulaMixer. The stained sample was incubated overnight at room temperature. Labeled and stained DNA was loaded onto the BioNano Genomics Saphyr chip. DNA was linearized in the nanochannel array by electrophoresis. Strands were imaged and the backbone and labels detected by BioNano image detection software. Single-molecule maps were assembled into consensus maps. Consensus maps were refined and merged based on overlapping segments. The final consensus maps were aligned to the GRCh38 human reference genome. Repeat expansions in the DMPK locus were identified by distances between flanking labels in the single-molecule maps aligned to the region.

### GRIP1 constructs and cloning

Double-stranded gene fragments encoding for GRIP1 kinesin-binding domain (GRIP1-KBD) +/- exon 21 were synthesized by Integrated DNA Technologies (IDT). The pEGFP-C1 (Clontech) plasmid containing a CMV promoter and C-terminal EGFP was used. Using the In-Fusion Directional Cloning Kit (Clontech), GRIP1-KBD +/- exon 21 was inserted into the pEGFP-C1 plasmid backbone between the SalI and BamHI sites. mCherry sequence was obtained from a pcDNA3.1 (Invitrogen) plasmid. An mCherry-C1 plasmid was subsequently constructed through excision and replacement of pEGFP between the AgeI and BglII sites. GRIP1-KBD −exon 21 was then cloned into mCherry-C1 as described above.

### Centrosome recruitment assay and quantification

Mouse Neuro2a cells^65^ were grown at 37°C in Dulbecco’s Modified Eagle Medium (Cytiva) containing 10% fetal bovine serum, D-glucose, L-glutamine, sodium pyruvate, streptomycin, and penicillin. Cells were trypsinized and plated onto glass chamber slides 1-2 days prior to transfection. To generate the BicD2-KIF5A expression plasmid, first a pcDNA3.1-BicD2-KIF1Balpha plasmid was generated. The pcDNA3.1-V5-His-TOPO backbone was cut with BamHI and EcoRV. BicD2 was amplified from a BicD2-FKBP expression plasmid provided by Gary Banker using primers GCTAGTTAAGCTTGGTACCGAGCTCGGATCCATGGATATCATGGATTACAAGGATGAC and CCACCCCCTCCCGAACCTCCGCCCCCTCTAGAGACGGTCCGATCT. The KIF1Balpha tail was amplified from mouse cDNA using primers GGAGGTTCGGGAGGGGGTGGCTCAGATACATCCATGGGGTCCCTC and GAGCGGCCGCCACTGTGCTGGATATCCTAGACTGTGGTTTCTCGACCT. The BicD2 PCR product and KIF1Balpha PCR products were joined together by PCR using the outer primers, and cloned into the pcDNA3.1 backbone by Infusion cloning (Clontech). The pcDNA3.1-BicD2-KIF1Balpha plasmid was then cut using SanDI and NotI, and the KIF5A tail was amplified from human cDNA using primers GCCTCAGTAAATTTGGAGTTGACTGC and TTAGCTGGCTGCTGTCTCTTGG, followed by GCTCAGATACATCCATGGGGTCCCTCGCCTCAGTAAATTTGGAGT and GGGCCCTCTAGACTCGAGCGGCCGCTTAGCTGGCTGCTGTCTCTT. The PCR product was cloned into the backbone also by Infusion cloning to generate the BicD2-KIF5A tail construct. Constructs containing FP-GRIP1-KBD +/- exon 21 and FLAG-BicD2-KIF5A were expressed by transfection using TransIT-LT1 (Mirus). Approximately 24 hours after transfection, cells were fixed using 4% paraformaldehyde and permeabilized with 0.2% Triton X-100. Cells were stained with a rabbit monoclonal anti-DYKDDDDK tag antibody (D6W5B; Cell Signaling Technologies) followed by an Alexa Fluor 647-conjugated goat anti-rabbit secondary antibody (Thermo-Fisher Scientific). Cells were then mounted in Fluoroshield (Millipore-Sigma). Cells were imaged using the Zeiss LSM880 microscope by epifluorescence with an AxioCam MRm camera and Apochromat 40x/1.3 NA objective. Tile scans were gathered for each condition using ZEN software (Carl Zeiss International). Image analysis and quantitation of tile scans were performed using ImageJ/Fiji. Intensity values for whole cells and centrosomes were manually traced and quantitated using EGFP or mCherry (whole cell) and FLAG (centrosome) signals as masks. Intensity values from these tracings were used to quantify recruitment of GRIP1-KBD to the centrosome by BicD2-KIF5A, as represented by the ratio of centrosomal signal to whole-cell signal.

## Supplemental Figures and Tables

**Figure S1.**
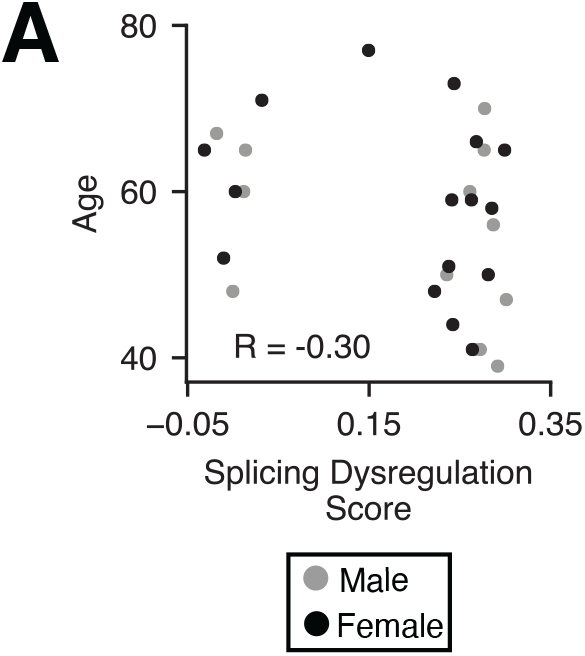
Splicing dysregulation does not correlate with age or show sex bias. A) Scatter of patient ages versus splicing dysregulation score with Pearson R = −0.30. Sex of each individual is indicated by differences in color; males are in gray and females in black.

**Figure S2.**
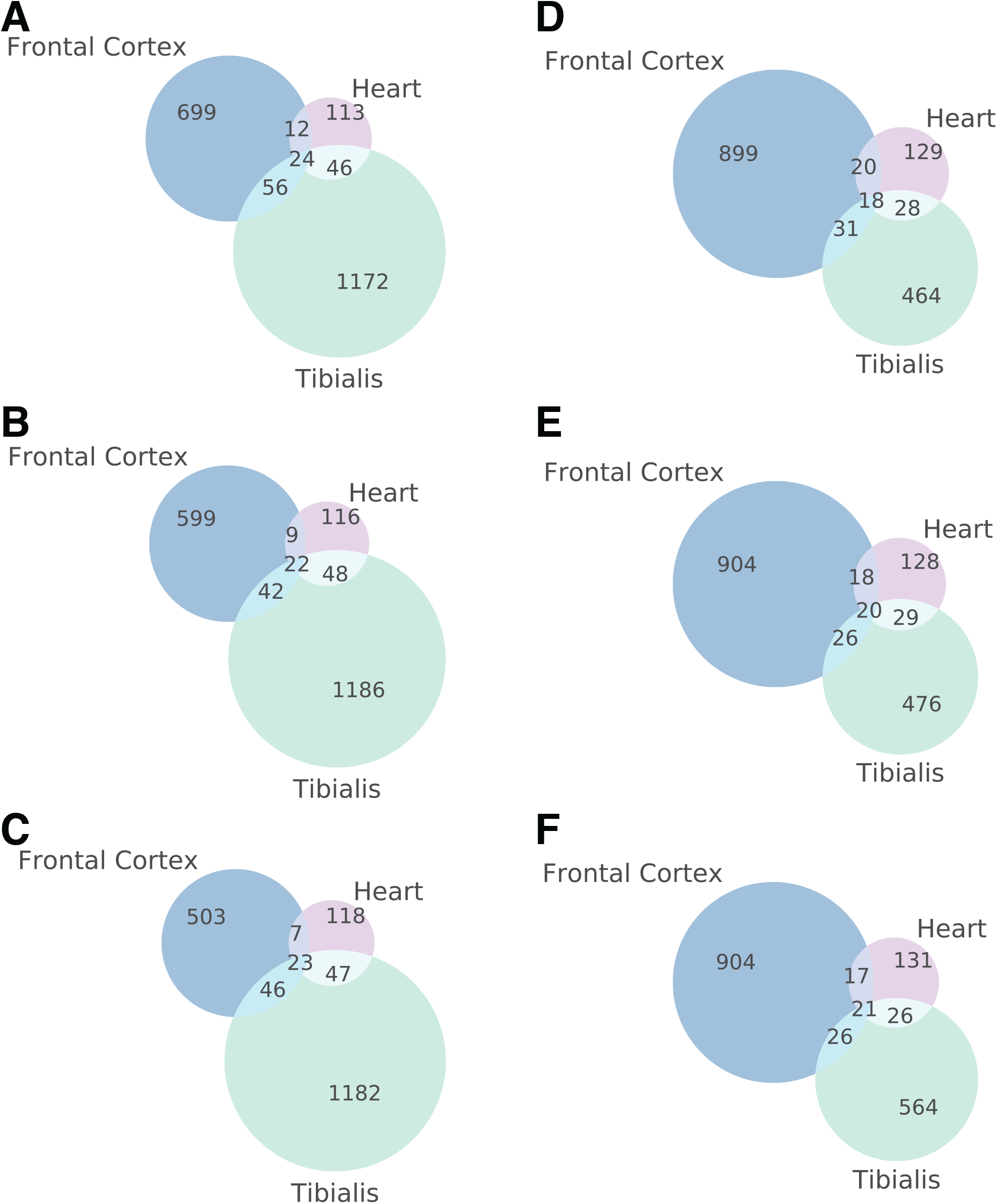
Patient cohort size largely affects the number of mis-spliced transcripts captured. A-C) Three separate random subsamples of the DM1 FC cohort were used to compute the number of exons dysregulated in FC, TA, and heart. 12 DM1 FC samples and 6 unaffected FC samples were selected in each subsample. All subsamples show that 15-19% of events significant in heart are also significant in FC, using |Δψ| ≥ 0.1 and p ≤ 0.01 (rank-sum test). Across the subsampling runs, the number of events identified to be significantly dysregulated in DM1 FC was 59-82% of the total 968 identified when using all samples, as shown in Figure 4A. D-F) Three separate random subsamples of the DM1 TA cohort were used to compute the number of exons dysregulated in FC, TA, and heart. All comparisons show that 23-26% of events significant in heart are also significant in TA, using |Δψ| ≥ 0.1, p ≤ 0.01 (ranksum test). Across the subsampling runs, the number of events identified to be significantly dysregulated in DM1 TA was 41-50% of the total 1298 identified when using all samples, as shown in Figure 4A.

**Figure S3.**
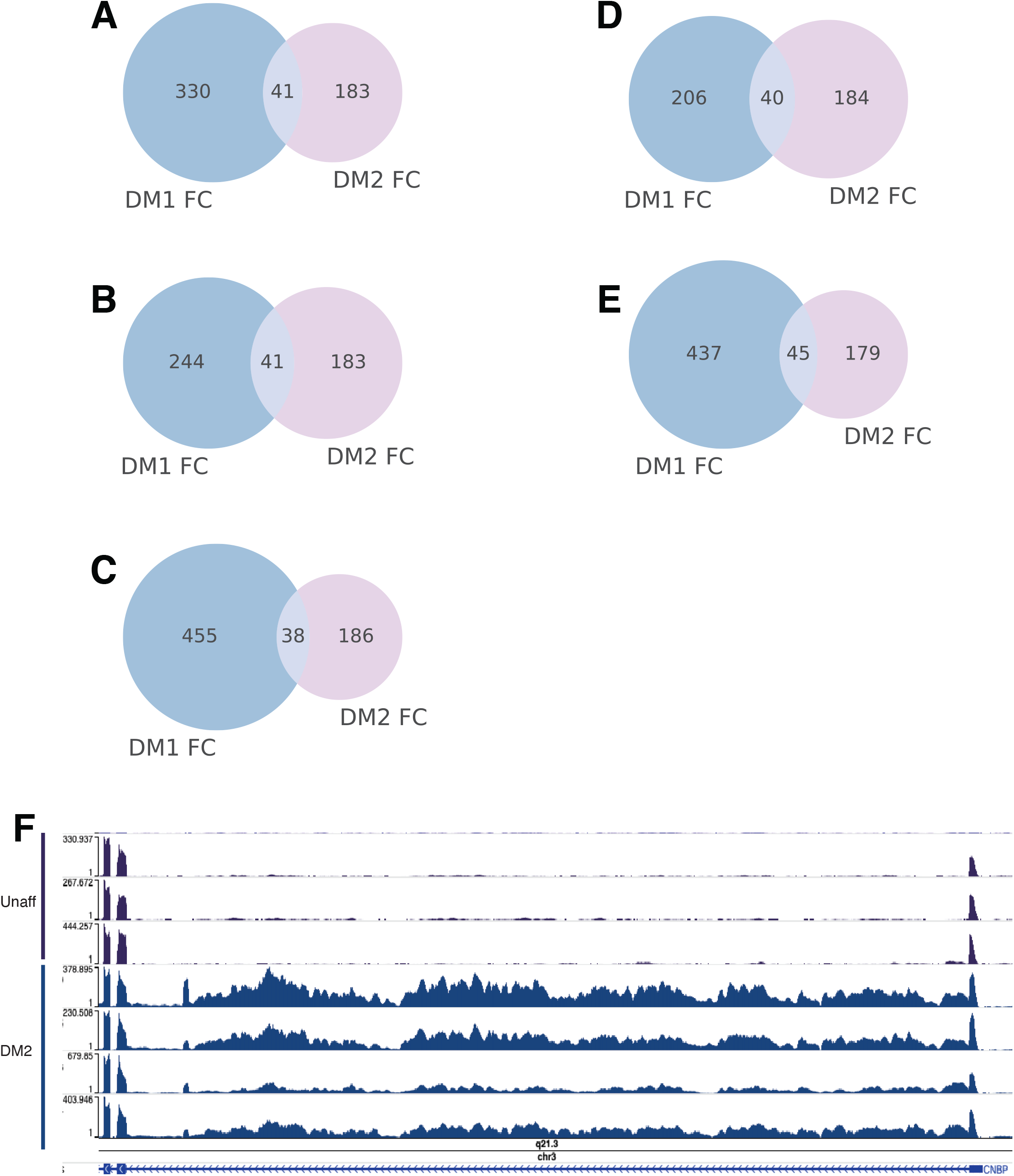
Small patient cohort size of DM2 limits global mis-splicing captured. A-C) Three separate random subsamples of the DM1 FC samples (4 DM1 patients, 8 unaffected controls) were used to compute the number of exons dysregulated in DM1, DM2, or both. All sub-samples show that 17-20% of the events dysregulated in DM1 were also dysregulated in DM2, using |Δψ| ? 0.1, p ≤ 0.01 (rank-sum test). The number of events significantly regulated in DM1 FC samples ranged between ~25-50% of the total 968 identified when using all samples, as shown in Figure 5A. D) Overlap of events when analyzing the 4 least severe DM1 patients (according to total splicing dysregulation) versus DM2 patients. E) Overlap of events when analyzing the 4 most severe DM1 patients (according to total splicing dysregulation) versus DM2 patients. F) Intron 1 retention in CNBP is observed in DM2 samples, but not in 3 representative unaffected controls.

**Figure S4.**
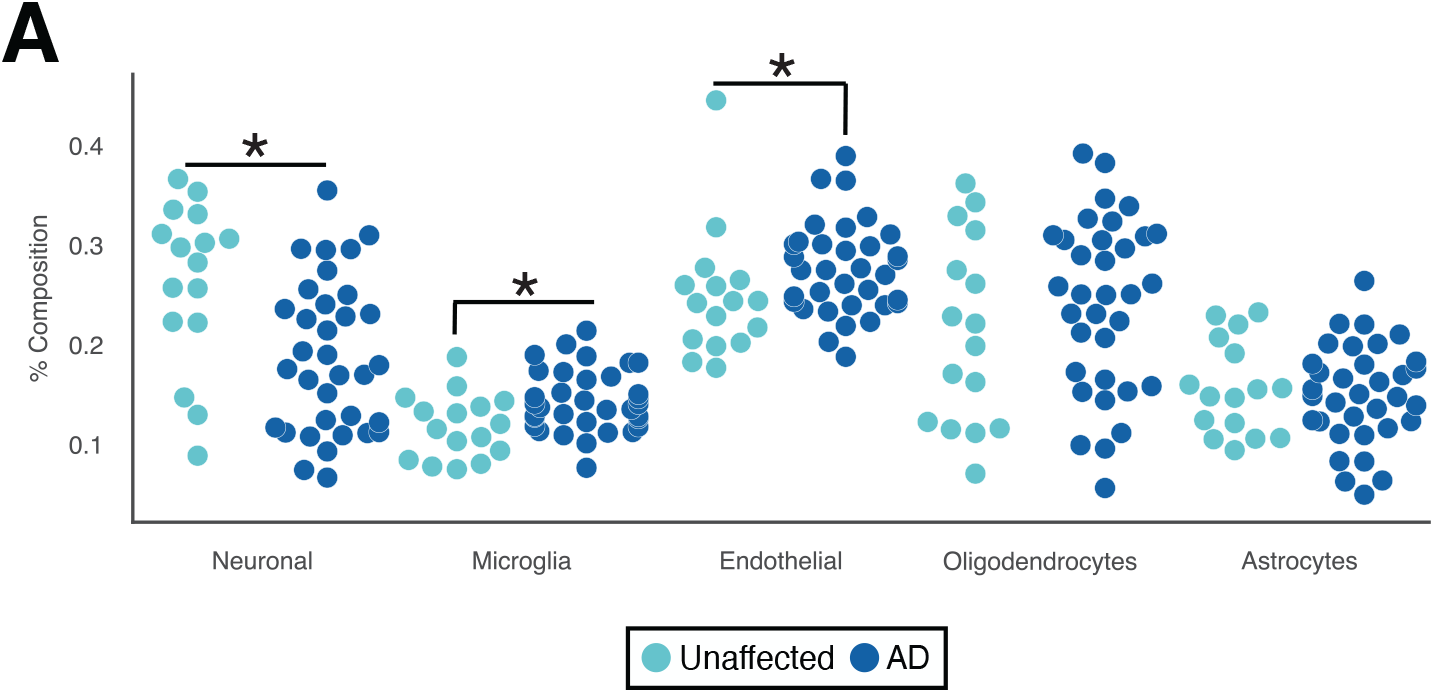
Cell composition analysis of the entorhinal cortex in AD patients shows significant neuronal loss, with increases in microglial and endothelial composition. A) RNA-seq data derived from 34 Alzheimer’s patients and 16 unaffected control entorhinal cortex samples were analyzed^45^. We estimated cell type composition using our Bayesian Inference approach. By rank-sum test, AD patients were found to have significantly lower neuronal composition and significantly higher microglial and endothelial cell composition.

**Table S1. Information about frontal cortex samples.** Column values are as follows: de-identified patient, disease status, age of patient at death, sex, RNA quality number (RQN), total RNA-seq reads across all runs performed, splicing dysregulation score, gene expression dysregulation score, 50^th^, 75^th^ and 90^th^ percentile of estimated number of CTGs captured in samples processed by optical mapping (BioNano).

**Table S2. Isoform estimates for DM1 and unaffected controls as assessed by MISO.** Column values are as follows: hg19 event coordinates, gene symbol, unaffected group mean percent spliced in (PSI, ψ), DM1 group mean ψ, ψ values for all 8 unaffected samples, ψ values for all 21 DM1 patients, whether the event was included in the top 130 most significant splicing events, whether the event was found to be significant in both frontal cortex and TA by |Δψ| ≥ 0.1, rank-sum p-value ≤ 0.01, whether the event was found to be significant in both frontal cortex and heart by |Δψ| ≥ 0.1, rank-sum p-value ≤ 0.01, and whether the event was found to be significant in both DM1 and DM2 by |Δψ| ≥ 0.1, rank-sum p-value ≤ 0.01.

**Table S3. Isoform estimates as assessed by MISO for orthologous exons in human and mouse.** Column values are as follows: hg19 event coordinates, mm10 event coordinates for orthologous exon, hg19 gene symbol, percent spliced in (PSI, ψ) values for all 8 unaffected humans, ψ values for all 21 DM1 patients, ψ values for all 3 MBNL DKO wild type (WT) mice, ψ values for all 3 MBNL DKO mice, ψ values for all 3 MBNL2 KO WT mice, and ψ values for all 3 MBNL2 KO mice.

**Table S4. Isoform estimates for DM2 and unaffected controls as assessed by MISO.** Column values are as follows: hg19 event coordinates, gene symbol, unaffected group mean percent spliced in (PSI, ψ), DM2 group mean ψ, ψ values for all 8 unaffected samples, ψ values for all 4 DM2 patients, whether the event was included in the top 59 most significant splicing events for DM2, and whether the event was found to be significant in both DM1 and DM2 by |Δψ| ≥ 0.1, rank-sum p-value ≤ 0.01.

**Table S5. Gene expression estimates for DM1 and unaffected controls as assessed by Kallisto.** Column values are as follows: hg19 gene symbol, gene expression values in TPMs for all 8 unaffected controls, gene expression values for all 21 DM1 patients, hg19 gene symbol, gene description, correlation of gene expression versus splicing dysregulation, whether this gene was found to be significantly regulated, whether it was up- or down-regulated, whether a protein product of this gene was found in CSF proteomics data, and whether protein products from this gene were found to have a signal peptide.

**Table S6. Gene expression estimates for DM2 and unaffected controls as assessed by Kallisto.** Column values are as follows: hg19 gene symbol, gene expression values in TPMs for all 8 unaffected controls, gene expression values for all 4 DM2 patients, hg19 gene symbol, and gene description.

